# Pronounced uptake and metabolism of organic substrates by diatoms revealed by pulse-labeling metabolomics

**DOI:** 10.1101/2021.11.04.467253

**Authors:** Nils Meyer, Aljoscha Rydzyk, Georg Pohnert

## Abstract

Diatoms contribute as a dominant group of microalgae to approximately 20% of the global carbon fixation. In the plankton, these photosynthetic algae are exposed to a plethora of metabolites, especially when competing algae are lysed. It is well established that diatoms can take up specific metabolites, such as vitamins, amino acids as nitrogen source, or dimethylsulfoniopropoionate to compensate for changes in water salinity. It is, however, unclear to which extent diatoms take up other organic resources and if these are incorporated into the cell’s metabolism. Here, we ask about the general scope of uptake of metabolites from competitors. Using labeled metabolites released during lysis of algae grown under a ^13^CO_2_ atmosphere, we show that the cosmopolitan diatom *Chaetoceros didymus* takes up organic substrates with little bias and remarkable efficiency. The newly developed pulse label/ mass spectrometry metabolomics approach reveals that polarity and molecular weight has no detectable influence on uptake efficiency. We also reveal that the taken-up pool of metabolites is partly maintained unaltered within the cells but is also subject to catabolic and anabolic transformation. One of the most dominant phytoplankton groups is thus substantially competing with other heterotrophs for organic material, suggesting that the observed absorbotrophy may substantially impact organic material fluxes in the oceans. Our findings call for the refinement of our understanding of competition in the plankton.

**Significance:** This study demonstrates a remarkably universal uptake of organic substrates by diatoms. The extent to which one of the most dominant phytoplankton groups is competing for organic material in the plankton is documented by novel pulse labeling metabolomics studies. Our results show that uptake of organic material by the photosynthetic microalgae occurs with remarkably little bias. Taken-up metabolites are further transformed by the diatoms or directly incorporated into the algal metabolome. Our study calls for a re-consideration of organic material fluxes in the oceans. Also, our understanding of competition in the plankton will have to be refined. The broader implications for the cycling of resources in plankton communities are discussed within this work.

## Introduction

The traditional view on marine plankton distinguishes between phytoplankton as primary producers and zooplankton as consumers (1). However, many planktonic eukaryotic organisms have been recognized as mixotrophs, which combine autotrophic photosynthesis with organic matter uptake (2). Many microzooplankton grazers are mixotrophic and retain functional algal organelles or even algal endosymbionts. Also photosynthetically active organisms, such as phytoflagellates and dinoflagellates can engulf and consume prey organisms to acquire nutrients (3). As an additional strategy, the uptake of dissolved organic carbon termed absorbotrophic mixotrophy or osmotrophy can be observed in microalgae. This process seems to be ubiquitous but clearly less understood (4).

First investigations of absorbotrophic mixotrophy in plankton focused on algae growth under extreme darkness in the presence of organic substrates (5, 6). Administration and uptake studies of radiolabeled substrates deepened our mechanistic understanding. However, experiments were always limited to investigating one compound or a compound class, such as specific vitamins or amino acids. It is now clear that absorbotrophic mixotrphy is widely distributed among planktonic eukaryotic organisms (7). However, the experiments with single compounds under limiting conditions conducted so far represent an oversimplification and do not reflect the situation in nature, where a cell is exposed to structurally most diverse metabolites. Consequently, the importance of absorbotrophic mixotrophy for pelagic food webs and for element cycling remains elusive and we are still far from quantitatively deciphering the trophic modes of phytoplankton (8).

The availability of organic substrates for uptake will be highly variable. In the plankton mass occurrences of algae, so-called algal blooms can last over days to weeks before the population breaks down and is succeeded by other species that become dominant. Especially during the decay of such algal blooms, the surviving competitors will be exposed to the metabolites of the lysed algae. Also, lysis of specific phytoplankton members by pathogens, such as algicidal bacteria or viruses results in situations where surviving resistant cells are exposed to the metabolomes of the lysed species.(9, 10) It is entirely unclear if and how all these compounds contribute to the metabolism of the phytoplankton and the potential ecological importance of phytoplankton as consumers of organic material is thus still poorly understood (11–14).

As critical primary producers, diatoms were initially classified as autotrophs, but the uptake capability of specific organic molecules was early recognized (14, 15). These include glucose, small polar organic acids such as acetate, succinate, fumarate, malate and lactate, amino acids, dipeptides and dimethylsulfoniopropionate (DMSP) (16–23). Most experiments provided mechanistic insight but did not accurately reflect natural conditions, where the water in which the algae live harbors a diverse mixture of organic compounds. Under natural conditions, cells are exposed to these metabolites and also light is available to support photosynthesis. Thus, two competing mechanisms for carbon acquisition, uptake and heterotrophy will be active. Here we address absorbotrophic mixotrophy in diatoms under non-limited conditions. The supply of nutrients and light in our study was non-limiting to allow efficient algal growth; organic metabolites were thus offered in addition to available inorganic sources.

We base our experimental setup on a well-investigated multi-partner interaction involving an algicidal bacterium and two cosmopolitan diatom species. *Kordia algicida* is a marine Flavobacterium that possesses algicidal activity leading to cell lysis of several microalgal species, including the diatom *Skeletonema costatum* (24, 25). *Chaetoceros didymus*, in contrast, is a naturally co-occurring diatom that is resistant against *K. algicida*. The impact of *K. algicida* was recently shown in field experiments where it induces a population shift in a natural phytoplankton community towards resistant algae (26). We hypothesize that during bacterial lysis of *S. costatum*, resistant species can benefit by taking up metabolites of the lysate in an absorbotrophic manner. Therefore, we exposed a culture of *C. didymus* to metabolites from a 50% diluted stationary culture of lysed *S. costatum* cells. We developed a novel analytical approach to test this hypothesis, including pulse labeling metabolomics and a novel non-discriminant data treatment routine. We show that uptake and metabolism of metabolites from the environment occur with remarkable efficiency in the resistant alga.

## Results

### Generation and evaluation of a labeled metabolome for uptake experiments

A complex medium containing the entire labeled metabolome of a diatom could be generated from ^13^C-labeled *S. costatum*. We grew *S. costatum* in a medium containing Na_2_^13^CO_3_ as a sole carbon source to obtain a labeled metabolome. With a repeated exchange of the medium, we reached up to 65% labeling in the algal metabolome. Mechanical lysis of the labeled culture and removing the cell debris gives an axenic medium, rich in organic metabolites (Suppl. Fig. 1).

We hypothesized that the resistant *C. didymus* encounters these metabolites also in the field when algicidal bacteria lyse its competitors. To test this hypothesis, labeled *S. costatum* cultures were infected with *K. algicida*, which resulted in lysis of more than half of the diatom cells within six days. Ultrahigh-pressure liquid chromatography-high resolution mass spectrometry (UHPLC-HRMS) of metabolites extracted from the medium revealed similar metabolic profiles in mechanically lysed *S. costatum* cells and those lysed by the bacteria (Table 1).

**Table 1:**
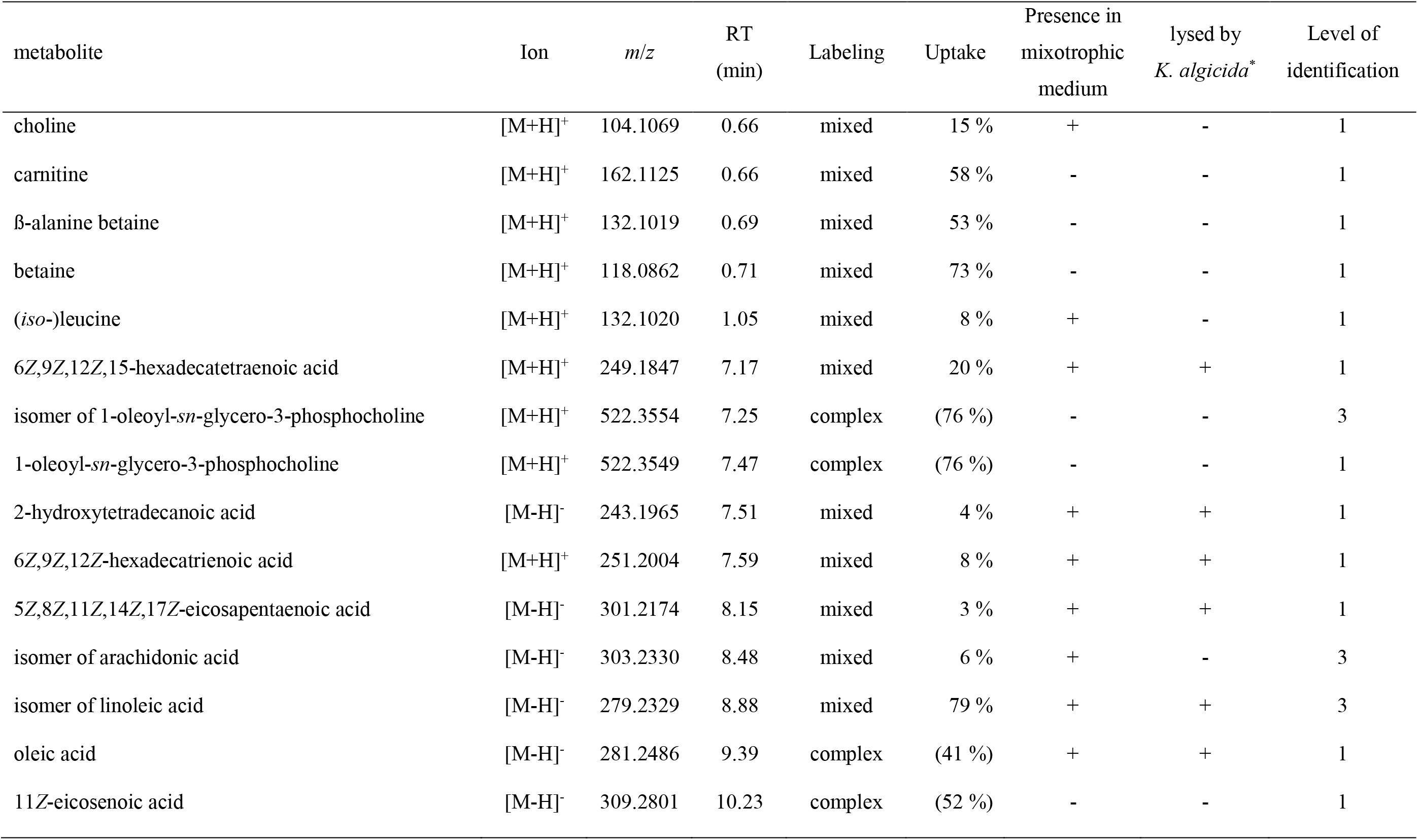
Labeled metabolites identified in the endometabolome of *Chaetoceros didymus*. *m*/*z*, mass to charge ratio; RT, retention time; labeling according to categories in Fig. 2; Uptake in % referring to the total amount of the respective metabolite; presence of the metabolite in the medium enriched with the *S. costatum* metabolome; presence in medium after lysis of *S. costatum* by *K. algicida*; level of identification according to Sumner, *et al.* (57), level 1 identified compound, level 3 putatively characterized compound class.

| metabolite | Ion | *m/z* | RT<br>(min) | Labeling | Uptake | Presence in<br>mixotrophic<br>medium | lysed by<br>*K. algicida* * | Level of<br>identification |
| --- | --- | --- | --- | --- | --- | --- | --- | --- |
| choline | [M+H]<sup>+</sup> | 104.1069 | 0.66 | mixed | 15 % | + | - | 1 |
| carnitine | [M+H]<sup>+</sup> | 162.1125 | 0.66 | mixed | 58 % | - | - | 1 |
| β-alanine betaine | [M+H]<sup>+</sup> | 132.1019 | 0.69 | mixed | 53 % | - | - | 1 |
| betaine | [M+H]<sup>+</sup> | 118.0862 | 0.71 | mixed | 73 % | - | - | 1 |
| (*iso*-)leucine | [M+H]<sup>+</sup> | 132.1020 | 1.05 | mixed | 8 % | + | - | 1 |
| 6Z,9Z,12Z,15-hexadecatetraenoic acid | [M+H]<sup>+</sup> | 249.1847 | 7.17 | mixed | 20 % | + | + | 1 |
| isomer of 1-oleoyl-*sn*-glycero-3-phosphocholine | [M+H]<sup>+</sup> | 522.3554 | 7.25 | complex | (76 %) | - | - | 3 |
| 1-oleoyl-*sn*-glycero-3-phosphocholine | [M+H]<sup>+</sup> | 522.3549 | 7.47 | complex | (76 %) | - | - | 1 |
| 2-hydroxytetradecanoic acid | [M-H]<sup>-</sup> | 243.1965 | 7.51 | mixed | 4 % | + | + | 1 |
| 6Z,9Z,12Z-hexadecatrienoic acid | [M+H]<sup>+</sup> | 251.2004 | 7.59 | mixed | 8 % | + | + | 1 |
| 5Z,8Z,11Z,14Z,17Z-eicosapentaenoic acid | [M-H]<sup>-</sup> | 301.2174 | 8.15 | mixed | 3 % | + | + | 1 |
| isomer of arachidonic acid | [M-H]<sup>-</sup> | 303.2330 | 8.48 | mixed | 6 % | + | - | 3 |
| isomer of linoleic acid | [M-H]<sup>-</sup> | 279.2329 | 8.88 | mixed | 79 % | + | + | 3 |
| oleic acid | [M-H]<sup>-</sup> | 281.2486 | 9.39 | complex | (41 %) | + | + | 1 |
| 11Z-eicosenoic acid | [M-H]<sup>-</sup> | 309.2801 | 10.23 | complex | (52 %) | - | - | 1 |

\*released from *S. costatum* during lysis by *K. algicida*

Before incubation with *C. didymus*, the metabolites from lysed *S. costatum* were sterilized and diluted with a medium containing inorganic carbon with normal isotope distribution (1.1% ^13^C, Suppl. Fig. 1). Thereby, we could ensure that ^13^C labeled organic metabolites taken up from the medium can be distinguished from *de novo* synthesized compounds. Following the same procedure, a control medium was generated using *S. costatum* cells grown in a medium with natural isotope distribution that could be used to generate mass spectra for structure elucidation. Using fragmentation trees, database comparison and subsequent comparison to authentic standards, we identified several of the labeled metabolites in *C. didymus* (Table 1).

### Evaluation of uptake

For uptake experiments, *C. didymus* was cultivated in a medium containing the sterilized labeled *S. costatum* metabolome or control medium for three days (n=3). We selected the concentration of added metabolites to be equivalent to 50% of those released by a lysed stationary culture. Cells were then recovered by filtration and washed extensively. UHPLC-HRMS analysis of the *C. didymus* metabolome, after being exposed to this medium under otherwise optimum growth conditions, revealed that the alga took up substantial amounts of labeled compounds from various metabolic classes. Quantitative analysis of labeling proved to be challenging in terms of chemoinformatic data treatment. Therefore, ions of the same metabolite, only differing in their number of incorporated ^13^C were summarized in an isotopologue group using the software X^13^CMS (27). Of 5587 isotopologue groups (positive and negative ionization mode) detectable in the endo metabolome 2381 were significantly labeled with ^13^C (Fig. 1A). After manual curation, 548 isotopologue groups were categorized according to their labeling pattern and analyzed regarding their retention time and mass to charge ratio. The degree of labeling was estimated using the probability mass function for Bernoulli trials (random experiments with precisely two possible outcomes, “success i.e. incorporation of ^13^C” and “failure i.e. incorporation of ^12^C “, in which the probability of success is the same every time the experiment is conducted):

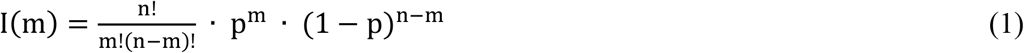

**Fig. 1:**
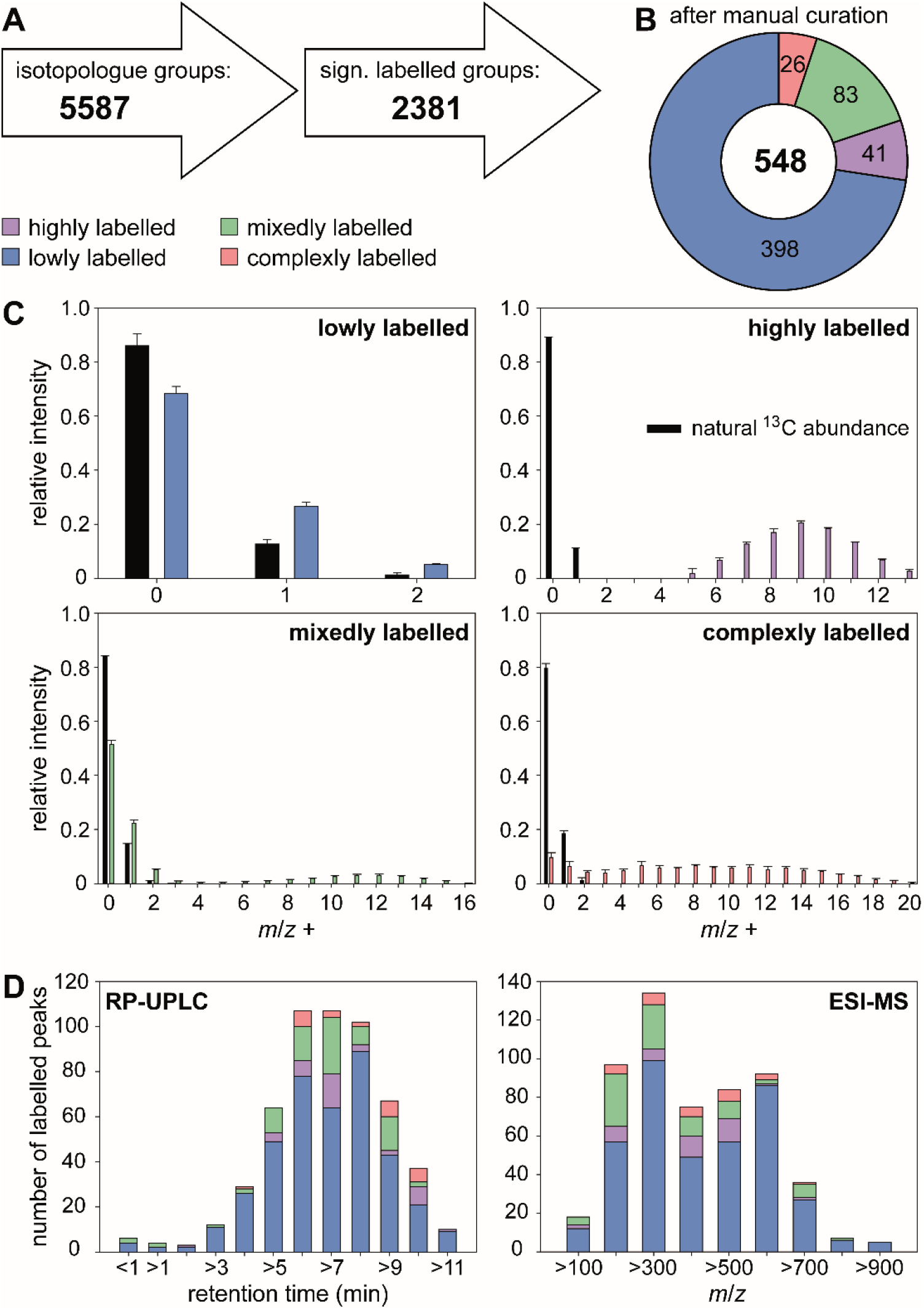
Labeling of metabolites in *Chaetoceros didymus* endometabolome after exposure to metabolites from lysed labeled *S. costatum*. (A) Isotopologue groups were detected and tested for significance using X^13^CMS. (B) After manual curation the remaining 548 isotopologue groups were categorized according to their labeling pattern. (C) Isotopologue distributions in metabolites with natural and enriched ^13^C abundance are depicted for each category. (D) Isotopologue groups were clustered by retention time and *m*/*z* range. A dot plot showing the correlation between retention time and *m*/*z* can be found in Suppl. Fig. 2.

The degree of labeling p for a metabolite with n carbon atoms is estimated from the intensities of a set of isotopologues I(m) with m ^13^C atoms (28).

### Classification of taken-up and processed metabolites

We categorized the metabolites according to their degree of labeling (Fig. 1B, C). 73% of the isotopologue groups were lowly labeled with a degree of labeling < 5% indicative of compounds mainly synthesized *de novo* in *C. didymus*. The low degree of labeling that still exceeds the natural ^13^C content of 1.1% can be explained by general utilization of taken-up metabolites in the metabolism: metabolites that are assimilated are catabolized to metabolic building blocks that are used together with the autotrophic metabolic pool and used for anabolism again. Seven percent of the isotopologue groups were highly labeled. They contained a degree of labeling similar to that of metabolites in the medium (ca. 65%). The cellular content of these highly labeled metabolites taken up from lysed *S. costatum* shows that *C. didymus* assimilates metabolites that it does not (or only to a minimal extent) produce itself. Certain compounds with a high degree of labeling can be found in *C. didymus* but not in the mixotrophic medium. These could arise from the metabolic transformation of more complex metabolites released by *S. costatum*. 15% of the isotopologue groups were labeled in a mixed manner. In the mass spectrum both, signals from a lowly labeled share and a highly labeled share of the respective compound can be detected. This pattern can be explained by metabolites that are biosynthesized by *C. didymus* and also acquired from the medium. This compound class includes a wide range of natural products, from small charged molecules like choline and carnitine to non-polar lipids and fatty acids. A few signals (5%) had a complex labeling pattern that Bernoulli statistics could not describe. These can be interpreted as compounds that result from metabolites taken up and further metabolized using the pool of *de novo* synthesized metabolites (Fig. 1B, C). For example, intermediate metabolic products like fatty acids can be utilized in the anabolism of more complex ones like lipids.

### Properties of taken up metabolites

Labeled metabolites span a wide range of polarity from charged small molecules to non-polar fatty acids. Most labeled products are found in the non-polar region of the chromatogram (Fig. 1D, Tab. 1, Suppl. Fig. 2). The identified polar metabolites that are efficiently taken up include the amino acids leucine and/or isoleucine and several small charged molecules, namely glycine betaine, β-alanine betaine, carnitine (58% of total cellular carnitine is labeled), and choline. Also, many fatty acids and lipids are taken up, including oleic acid, 5*Z*,8*Z*,11*Z*,14*Z*,17*Z*-eicosapentaenoic acid, 6*Z*,9*Z*,12*Z*-hexadecatrienoic acid, 6*Z*,9*Z*,12*Z*,15-hexadecatetraenoic acid, 2-hydroxytetradecanoic acid and isomers of linoleic and arachidonic acid as well as the lipid 1-oleoyl-*sn*-glycero-3-phosphocholine.

Metabolites that were taken up also span a wide *m*/*z* range, with a maximum between *m*/*z* 200 and *m*/*z* 700 (Fig. 1D, Tab. 1, Suppl. Fig. 2). There is thus relatively little size discrimination for the uptake.

### Analysis of selected metabolites

Detailed analyses of the isotopic pattern enabled us to determine the ratio of heterotrophic uptake to *de novo* biosynthesis and look for evidence of mixed strategies. For example, the isotopologues of 6*Z*,9*Z*,12*Z*,15-hexadecatetraenoic acid (Fig. 2A) originate from two distinct pools (Fig. 2D), a lowly labeled pool from *de novo* biosynthesis and a highly labeled pool from uptake. Modeling with Bernoulli statistics showed that the *de novo* biosynthesis pool had a degree of labeling of 2.8%, slightly higher than what would be expected from photosynthesis using exclusively natural inorganic carbon (Fig. 2B). The highly labeled pool has a very similar isotopologue distribution compared to the fatty acid from the lysed labeled alga (Fig. 2C). Modeling a mixture of these two pools showed that ca. 20% of the metabolite in the algae results from uptake, while 80 % are synthesized de novo. This complementation of de novo synthesized products with externally available metabolites is observed for several metabolites with variable proportions of the two sources from only a few percent to nearly 80% of the metabolite acquired by uptake (Table 1).

**Fig. 2:**
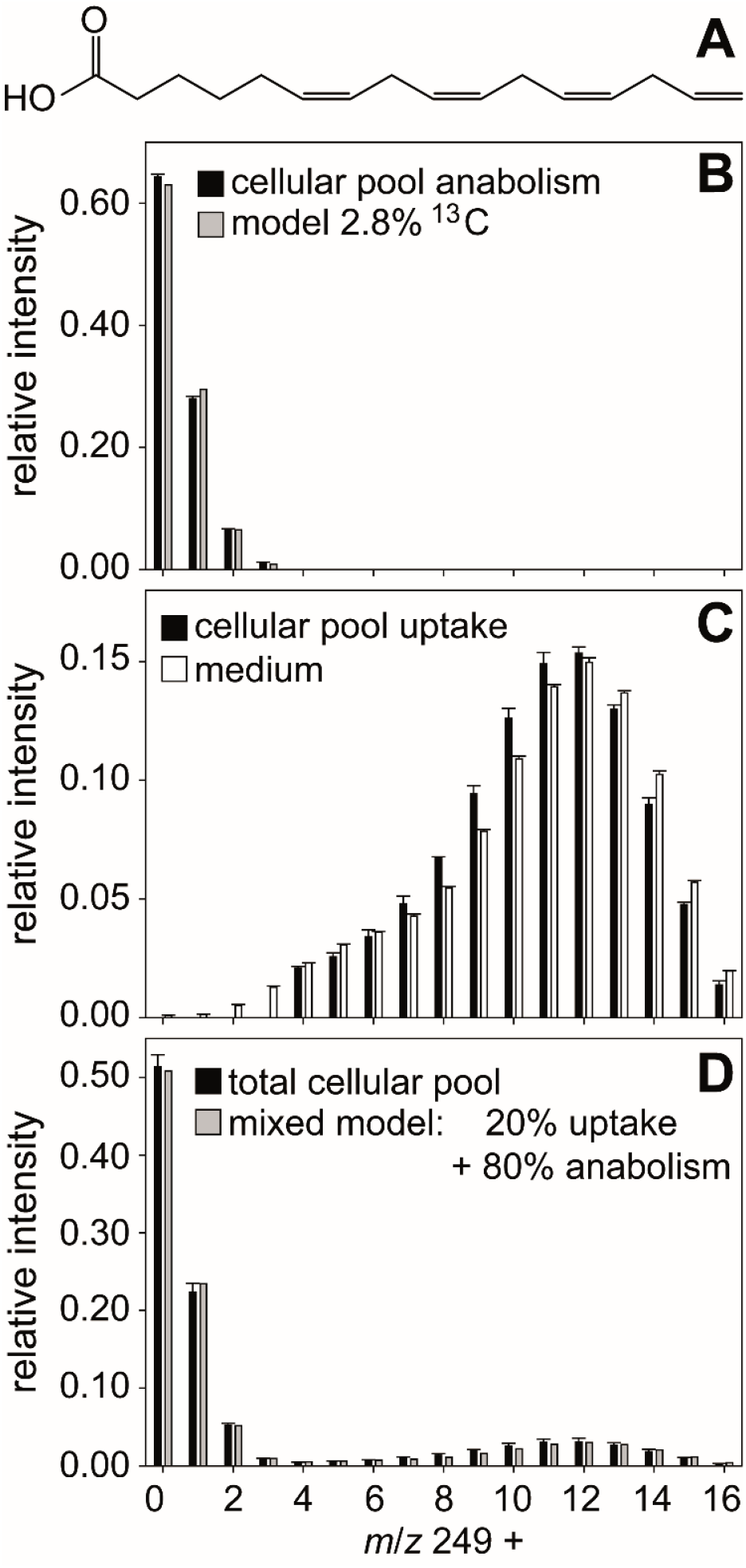
Labeling pattern of hexadecatetraenoic acid. (A) A mixed labeled metabolite was identified as 6*Z*,9*Z*,12*Z*,15-hexadecatetraenoic acid. For data evaluation the measured mass spectrum (black bars in D) was divided into a lowly labeled (B) and a highly labeled pool (C). Modelling (grey bars) shows that the lowly labeled pool contains 2.8 % ^13^C and thus likely derives from anabolism. (C) The highly labeled pool is taken up from the medium, black bars represent measured data of cellular hexadecatetraenoic acid, white bars measured data of hexadecatetraenoic acid in medium. (D) The measured mass spectrum in D can be explained by 20% hexadecatetraenoic acid derived from uptake and 80% from *de novo* synthesis (grey bars). All data are mean ± SD from biological triplicates.

The isotopologue patterns of metabolites that can be explained by an uptake of resources from the medium followed by transformations within the cell using the pool of *de novo* synthesized precursors are more complex. They do not follow the Bernoulli statistics since different resources can be utilized in different relative amounts. An example is the lysophosphatidylcholine shown in Fig. 3A. The isotope pattern of the lipid (Fig. 3B) cannot be interpreted with the model described above, but tandem MS experiments allow to dissecting the lipid. This reveals a unique labeling pattern for those parts of the molecule that are derived from different biosynthetic pathways. The isotopologue pattern of oleic acid in the lysophosphatidylcholine (Fig. 3C) can also not be fitted with a Bernoulli statistic. The fatty acid is thus assembled using resources that were taken up as well as *de novo* produced. This labeling pattern in the fatty acid moiety is also observed in free oleic acid and 11*Z*-eicosenoic acid. Lipid assembly thus does not discriminate between acquired and *de novo* synthesized resources. In contrast, the choline fragment detected in the same substance shows that most choline is highly labeled and therefore taken up (Fig. 3D). The glycerol moiety of the molecule is not giving charged fragments, but its labeling could be established indirectly. Therefore, we conducted fragmentation of the M+8 isotopologue of the lysophosphatidylcholine. The MS/MS of this ion gave rise to an oleic acid fragment with isotopologues containing down to zero ^13^C (Fig. 3E). The remaining eight carbon atoms in the uncharged C8-fragment have thus to be labeled in different degrees. The fragments can thus only be derived from a precursor with labeled, thus acquired glycerol.

**Fig. 3:**
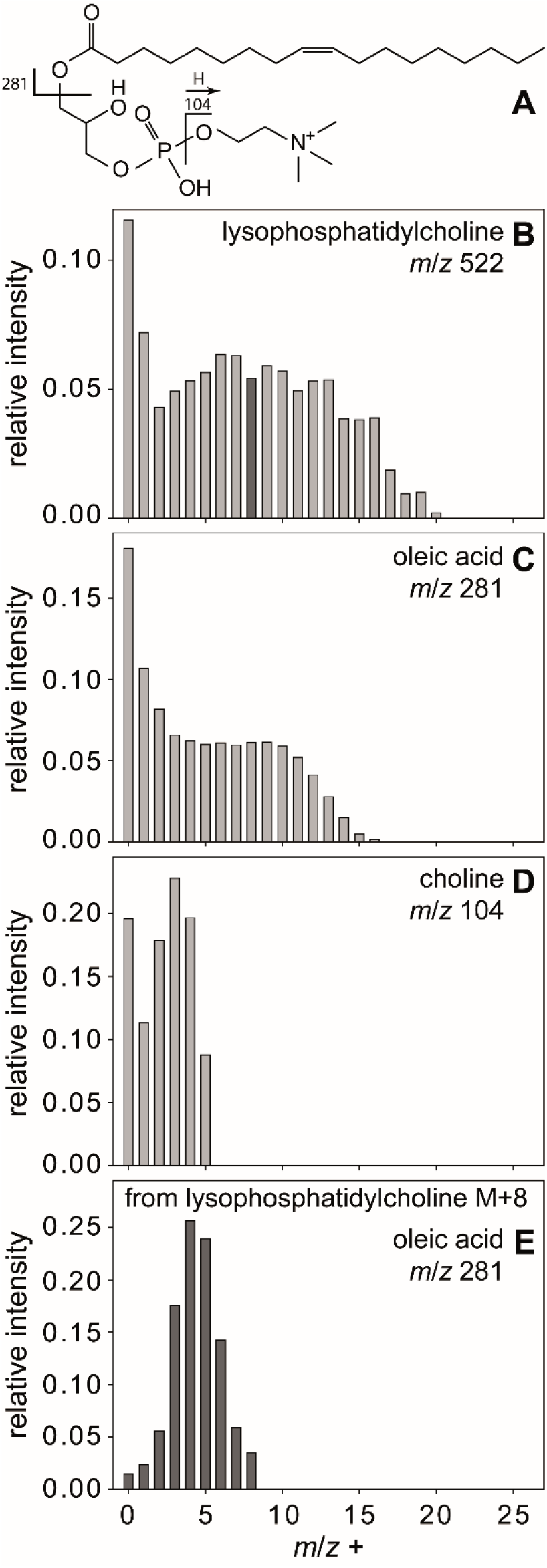
Complex labeling of lysophosphatidylcholine. All isotopologues (B) of lysophosphatidylcholine (A) are fragmented to yield labeling of the building blocks oleic acid and choline (D). Fragmentation of lysophosphatidylcholine M+8 isotopologue (black bar in B) yields oleic acid (E) with less than four ^13^C, thereby indirectly proving the labeling of glycerol.

These exemplarily discussed mass spectra stand for several hundreds of labeled peaks in the chromatogram of the metabolome of exposed *C. didymus* (see Fig. 1 and Supporting information). Using the combined results, we can now draw a picture of the absorbotrophic metabolism in *C. didymus* (Fig. 4). The uptake introduced here for *C. didymus* is not limited to this one species but broader distributed in diatoms. When the diatom *Thalassiosira weissflogii* was raised in the labeled medium as described for *C. didymus*, labeling patterns similar to the ones described above were detected (data not shown).

**Fig. 4:**
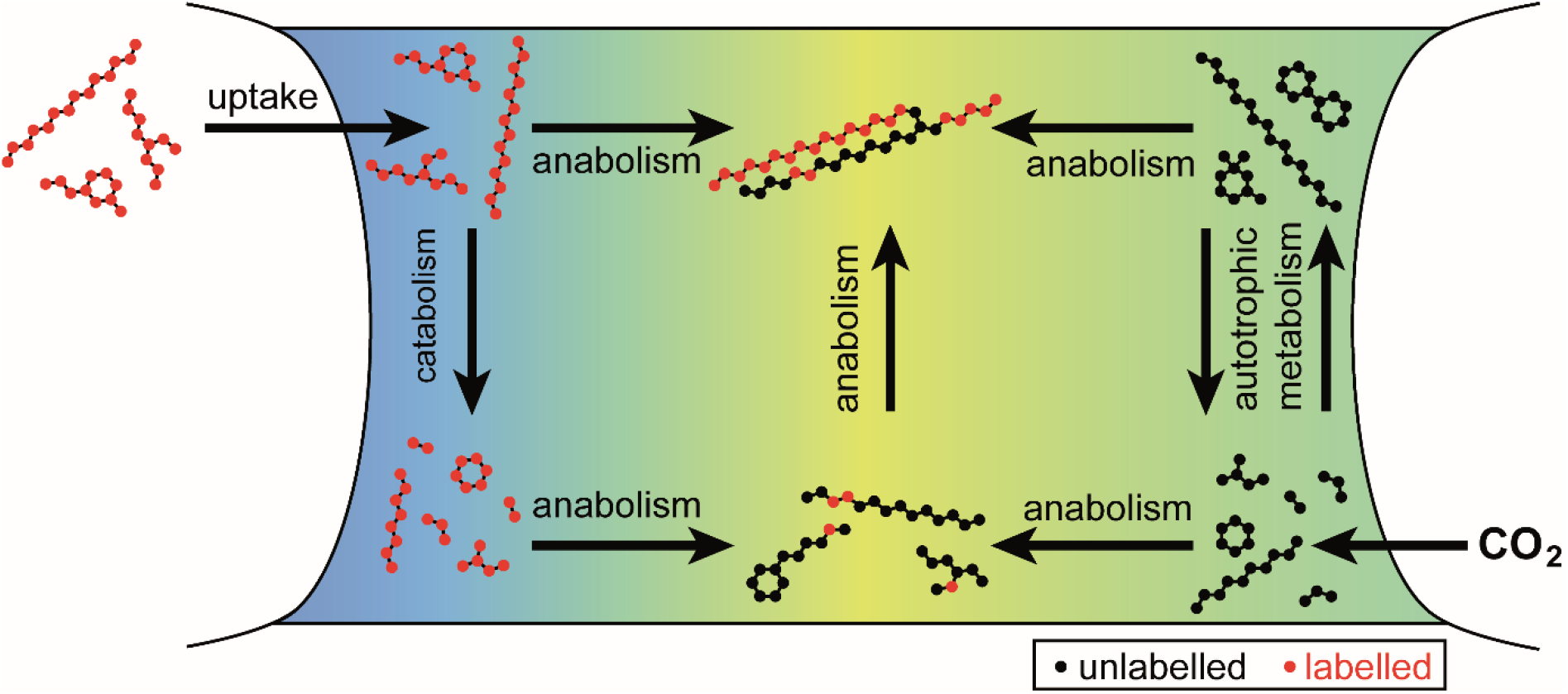
Uptake and metabolism of labeled organic compounds by *Chaetoceros didymus*. Organic compounds labeled with ^13^C (red) are taken up and transformed. Labeled compounds and their catabolic products are mixed with unlabeled metabolites (black) from autotrophic metabolism in anabolic reactions.

## Discussion

### ^13^C labeling allows tracing of metabolic shuttling between microalgae

In order to study potential absorbotrophic mixotrophy in the diatom *C. didymus* a medium rich in ^13^C labeled organic metabolites was prepared by mechanical lysis of a globally labeled *S. costatum* culture and removal of the cell debris by filtration. The resulting metabolome enriched medium showed a similar (but labeled) metabolic profile as the medium of an infection experiment where the lytic bacterium *K. algicida* lysed *S. costatum*. We thus conclude that *K. algicida* is a sloppy feeder not utilizing the entire algal metabolome but instead leaving substantial organic resources that it does not require or recover in the water. Surviving algae, such as the resistant competitor *C. didymus* will be exposed to these metabolites. Earlier, we observed that such exposure to metabolites released from lysed algae supports the growth of *C. didymus* if administered at low concentrations. However, no information about the underlying mechanism was available (29). The ability to take up released organic molecules may counterbalance the metabolic costs to maintain resistance mechanisms and would be highly advantageous, for example, during the collapse of a competing algal bloom as it was simulated in this study (10, 30). If the metabolic uptake also compensates in phases of very dilute phytoplankton abundance would have to be verified in follow-up studies. We have, however, no indication from the analysis of labeling that there would be a bias for the uptake of higher concentrated metabolites.

We reached up to 65% labeling in the *S. costatum* metabolome using repeated exchange of medium containing Na_2_^13^CO_3_ as an exclusive carbon source. Analysis of the *C. didymus* endometabolome after being exposed to the labeled metabolome of *S. costatum* for 3 days under otherwise optimum growth conditions revealed that the alga took up substantial amounts of labeled compounds from various metabolic classes. Given that analysis of every single isotopologue of a labeled metabolite would generate highly complex and partially redundant information, we reverted to a statistical treatment assuming that labeling results in an isotopologue distribution that can be described with a Bernoulli statistic. Indeed, this data treatment allowed to match mass spectral patterns of labeled metabolites to predicted spectra that would result from specific degrees of labeling. Thereby the average 65% labeling of the metabolome could be determined.

Labeling patterns of metabolites in *C. didymus* after exposure to ^13^labeled *S. costatum* medium

Analysis of the metabolome of the resistant alga *C. didymus* after exposure to ^13^C-labeled *S. costatum* revealed unlabeled metabolites, metabolites with unaltered full labeling and those with more complex mass spectra that could be assigned to different metabolic processing (Fig. 1). The methodology described here thus not only shows the uptake of one metabolite, but allows simultaneous quantification of the uptake and analysis of the metabolic fate of all taken up metabolites in the receiving alga. This is a valuable expansion of the classical fluxomics approach, where feeding of one single labeled metabolite to heterotrophs can only reveal its uptake and metabolism. Our experimental approach reflects the situation in the plakton with highly complex microbial communities and complex exometabolomes (31). It allows evaluation of the uptake capacity and incorporation of the taken-up metabolites under ecologically relevant conditions.

### General patterns in the uptake of organic metabolites by the receiving diatom C. didymus

Detailed analysis of the mass spectra allows to draw a picture of absorbotrophy in *C. didymus* (Fig. 4). More than a quarter of all detected metabolites in *C. didymus* were labeled to different degrees (Figure 1). Nearly 10 % showed the identical labeling pattern as those in the ^13^C-labeled *S. costatum* metabolome. These metabolites are not (or only to a minor extent) synthesized by the receiver but taken up and maintained in the cells. The major part of taken-up metabolites showed mixed labeling, indicative for the uptake of a metabolite that is also synthesized by the receiver. This includes a wide range of natural products from small charged molecules like choline and carnitine to non-polar lipids and fatty acids. Metabolites acquired from the outside water can thus be metabolized in the same way as the de novo produced compounds, indicating that no compartmentation of assimilated material occurs. Instead, the autotrophic and the phototrophic pool are used for anabolism and catabolism. For example, the presence of complex labeling patterns can be explained by the use of intermediate fatty acids in the anabolism of more complex lipids. The complex isotopologue pattern of oleic acid and 11*Z*-eiosenoic acid, for example, might reflect a dynamic system with rounds of beta oxidation releasing labeled acetate and subsequent fatty acid re-assembly from this labeled and the own unlabeled acetate pool. Since the isotopologues of these fatty acids are not Bernoulli-distributed the relative amount of labeled acetate in the total acetate pool used for fatty acid biosynthesis might vary over time.

### Utilization of assimilated metabolites

The fact that uptake has no apparent preference for nitrogen containing metabolites contradicts the assumption that diatoms use specific heterotrophic mechanisms to acquire reduced nitrogen (19). Notably, many metabolites with essential physiological functions are thus not exclusively produced *de novo* but taken up in high proportions. Control of physiological concentrations will thus have to include biosynthesis, catabolism as well as uptake. A universal uptake could be highly advantageous in the event of a lysis of a competing algal bloom (10, 30). But even under regular conditions in plankton, algae might encounter metabolites released by other members of the phytoplankton that they can take up and benefit from (32).

The observation of universal uptake of metabolites requires re-thinking of the interpretation of incubation studies with single labeled substrates. It is well documented that specific primary metabolites can be taken up by the cells under limiting conditions. Thus, under darkness, uptake of organic acids like lactate and malate was observed (21). Amino acid assimilation after peptide lysis was interpreted as nitrogen uptake mechanism (19) and uptake of the essential metabolite dimethylsulfoniopropionate (DMSP) was discussed as a way to compensate for the lack of own DMSP biosynthetic capabilities (23). We now expand this view and introduce that diatoms can take up a structural variety of metabolites released from competitors even under optimized growth conditions. Thus no specific compensation mechanisms but rather universal complementation of the own metabolome occurs but rather an universal supply with potentially valuable metabolites. This unspecific uptake of metabolites over a wide range of polarities and masses newly defines diatoms as a general sink of organic carbon in the sea.

### Physiological function of assimilated metabolites

Of the multiple labeled metabolites, two groups will be exemplarily highlighted here, polar nitrogen containing metabolites and fatty acids. Polar metabolites that are taken up include glycine betaine, β-alanine betaine, carnitine and choline, with diverse physiological functions. Glycine betaine is a known osmolyte and uptake in diatoms has already been described under N-limited conditions (33, 34). Choline is a biosynthetic precursor of glycine betaine (35) and important as structural element in phospholipids (11). It has been found as a free form and in lysophosphatidylcholine in this study (Table 1). However, the labeling pattern of the choline fragment of lysophosphatidylcholine differs from the one of the free form (Suppl. Fig. 3), indicating different origins. β-alanine betaine is a known osmoprotectant in plants and is biosynthesized via the methylation of β-alanine, a building block of coenzyme A (36). It has also been found in marine algae (37). Carnitine is a central metabolite in energy metabolism of all eukaryotic cells. It plays an essential role for the transport of fatty acids across the mitochondrial membrane. The utilization of carnitine by diatoms has been reported previously by measuring oxidation rates in a biofilm-forming freshwater diatom (38). The presence of labeled *N*-methyl groups (Suppl. Fig. 4) excludes lysine degradation (39) as a metabolic origin. The high proportion of heterotrophically acquired carnitine and the presence of many fatty acids among the taken-up metabolites is striking. It suggests a highly active transport mechanism for these high-energy fatty acids and the *N*-containing carnitine. It would be interesting to investigate whether an acylcarnitine-type transport system might facilitate the uptake of fatty acids across the cellular membrane. The presence of a protein homologous to a class I carnitine/acylcarnitine translocase in a diatom cell wall proteome supports this hypothesis (40). Several labeled fatty acids and derivatives have also been identified (Table 1). Eicosapentaenoic acid is one of the dominant fatty acids in diatoms and precursor of many bioactive oxylipins (11, 41–43). 6*Z*,9*Z*,12*Z*-hexadecatrienoic acid is the precursor of octadienal, an allelopathic polyunsaturated aldehyde (44). Also, oleic acid, 11*Z*-eicosenoic acid, and 6*Z*,9*Z*,12*Z*,15-hexadecatetraenoic acid are common in diatoms (43, 45–48). Two other C18- and C20 polyunsaturated fatty acids were identified according to retention time and mass spectra. Comparison with synthetic standards showed that these fatty acids were isomers of linoleic and arachidonic acid. Fatty acids and their derivatives have multiple physiological functions. The fact that they are present in labeled and unlabeled forms in after the incubation experiments indicates an unbiased utilization of de novo and assimilated compounds. This might serve as a mechanism to avoid costly biosynthesis of fatty acids, which is supported by the observation that the fatty acids are not only internalized or adsorbed to lipidic structures due to their physicochemical properties but rather incorporated into the primary metabolism of the diatom.

The broad range of polarity and molecular weight of assimilated compounds raises the question of the uptake mechanism. Candidate systems for fatty acid shuttling include specific transporters and genome data shows homolog candidate sequences (40, 49). But indeed, also unspecific absorption and incorporation mechanisms that do not require transporters will most likely be involved. Nevertheless, the question of how diatoms acquire exogenous metabolites is still open.

### Concluding remarks

The diatom *C. didymus* takes up and incorporates metabolites from lysed competitors with surprising little bias. The same is true for another model diatom *T. weissflogii*, which might suggest a rather universal mechanism. The multitude of cellular functions in which these metabolites are involved suggests that uptake complements the internal metabolic pool resulting from many different biosynthetic pathways.

The universal absorbotrophic lifestyle of one of the most abundant algal classes in the oceans substantially changes our view of the metabolic shuttling in phytoplankton communities. These algae take up not only a few highly polar metabolites for e.g. nitrogen supply. They rather complement their metabolism quite universally with resources from the surrounding seawater. This process occurs even under illumination and is not related to the complementation of lacking photosynthate in the dark. Thus, in addition to bacteria, diatoms compete for the dissolved organic carbon in the plankton (50, 51). Our study has consequences for element cycling in the oceans and ecosystem dynamics that will have to be addressed in the future.

## Materials and Methods

### Experimental design

Diatoms were grown under optimum conditions and exposed to a labeled metabolome of a competitor. The diatoms were extracted using optimized protocols for metabolomics sampling and extracts were analysed by liquid chromatography mass spectrometry to evaluate the uptake of labeled metabolites.

### Algal culturing

*C. didymus* was isolated by W. Kooistra from the Gulf of Naples (Stazione Zoologica Anton Dohrn, Naples, Italy) and *S. costatum* was obtained from the Roscoff Culture Collection (Roscoff, France). Both algae were cultivated in batch culture using artificial seawater medium(52) in 50 mL Greiner Bio-One cell culture flasks at 11-13°C under a 14:10 h light: dark regime with an illumination of 20-25 μmol photons m^−2^ s^−1^. Development of cultures was followed by *in-vivo* Chl *a* fluorescence using a Mithras LB 940 plate reader (excitation 430 nm, emission 665 nm).

### Global ^13^C-labeling

For global ^13^C-labeling of *S. costatum* we used autoclaved artificial seawater medium that was prepared without addition of NaHCO3. An aliquot of this medium was utilized to dissolve NaH^13^CO_3_ (98 atom %, Sigma-Aldrich, Munich, Germany). This solution, containing sufficient NaH^13^CO_3_ to reach a final concentration of 2.38 mM, was sterile filtered (0.2 μm pore size, Sarstedt Filtropur S) and transferred back to the medium bottle. Tissue culture flasks were filled to the neck in order to minimize the area for CO2 exchange with the atmosphere and were inoculated with < 1 % (v/v) of a stationary *S. costatum* culture. After growing to stationary phase, an aliquot was taken and transferred to fresh ^13^C-enriched medium (< 1 % (v/v)). After two of these cycles a plateau in the degree of labeling (verified by mass spectrometry as described below) was reached and the cultures were used for further experiments.

### Bacterial culturing

*K. algicida* (30) was cultivated on marine broth agar at 30°C for 2 days. The bacterial lawn was removed with a sterile cotton swab and re-suspended in algal culturing medium to an OD550 of 0.5 determined on a Genesys 10S UV-Vis spectrophotometer (Thermo Fisher Scientific, Waltham, MA, USA).

### Co-culturing experiment and extraction of released metabolites

*S. costatum* cultures reared in ^12^C or ^13^C medium were co-cultured in triplicates (185 mL each) with the *K. algicida* added to a final OD550 of 0.01. After 6 days the lysed cultures were gently filtered (0.2 μm pore size) and the flow-through extracted for metabolome analysis as follows. Solid phase extraction cartridges (Chromabond easy, Macherey-Nagel, Düren, Germany) were equilibrated with 4 mL of methanol (Chromasolv^©^ Plus, Sigma-Aldrich, Munich, Germany) and 4 mL of water (Chromasolv^©^ Plus, Sigma-Aldrich, Munich, Germany) before the filtrate (170 mL) was applied using vacuum with a flow rate < 1 L h^−1^. The cartridge was washed with 4 mL water, air-dried and then extracted via gravity flow using 2 mL of methanol followed by 2 mL of methanol/tetrahydrofuran 1:1 (tetrahydrofuran HiPerSolv, VWR, Dresden, Germany). This extract was frozen until further chemical analysis.

### Tests for mixotrophy

Stationary cultures (45 mL) of *S. costatum* reared in ^12^C or ^13^C medium were centrifuged (500 x g, 15 min, 10°C) and washed three times by repeated addition of 45 mL of ^12^C medium to the harvested pellets and centrifugation. After the third washing step the supernatant was collected as control medium (this was processed in parallel to the cells and later served as control for the natural ^13^CO_2_) and the cell pellet was re-suspended in 45 mL ^12^C medium. To disrupt cells, the suspension was frozen at −20°C, thawed and treated in an ice-cold ultrasonic bath for 10 min. The lysate was filtered (1.2 μm pore size, GF/C, Whatman, GE Healthcare, Little Chalfont, United Kingdom), acidified to pH ≤ 1 for sterilization using 30% hydrochloric acid, incubated at 0°C for 10 min and subsequently neutralized under sterile conditions using a saturated sodium hydroxide solution. The solutions were stored at −20°C until use. After thawing, the solutions were diluted 1:1 with ^12^C medium to yield the final medium for the investigation of mixotrophy (^12^C organic & ^12^C inorganic or ^13^C organic & ^12^C inorganic). 45 mL aliquots of both media were extracted as described above for exometabolomic analysis to determine the organic metabolites. For determination of mixotrophy 110 mL aliquots of these media were inoculated with 1% (v/v) stationary *C. didymus* culture in triplicates and cultivated for 3 days. Directly after inoculation and after 3 days of cultivation, samples (45 mL each) for intra- and extracellular metabolomics were processed. Samples were filtered (1.2 μm, GF/C, Whatman, GE Healthcare, Little Chalfont, United Kingdom) and the flow-through processed for exometabolomic analysis as described above (see co-culturing experiment). The cells were washed off the filter with an ice-cold freshly prepared mixture of methanol/ethanol/chloroform (1:3:1) (ethanol LiChroSolv^©^, Merck, Darmstadt, Germany; chloroform HiPerSolv^©^, VWR, Dresden, Germany). Extracts were treated in an ultrasonic bath for 10 min, centrifuged (30,000 x g, 15 min, 4°C) and the supernatant was stored at −20°C. As a control to prove that no NaH^13^CO_3_ enrichment was present in mixotrophic media, the third wash supernatant (see above) was used as medium to cultivate *C. didymus* and metabolites were extracted as described. A graphical representation of the experimental setup can be found in Suppl. Fig. 1.

### Analysis of exo- and endometabolomes with LC-MS

Extracts from cells and media (see above) were dried in a nitrogen flow at room temperature and were resuspended in up to 200 μL methanol. Metabolites were separated on an UltiMate 3000 UHPLC (Thermo Fisher Scientific, Waltham, MA, USA) equipped with an Accucore C18 column (100×2.1 mm, 2.6 μm) at 25°C using water with 2% acetonitrile and 0.1% formic acid (A) and pure acetonitrile (B) as mobile phase. The gradient was as follows: 100% A for 0.2 min, linear gradient to 100% B in 7.8 min, 100% B for 3 min, linear gradient to 100% A in 0.1 min, 100% A for 0.9 min. The UHPLC was connected to a QEplus Orbitrap mass spectrometer (Thermo Fisher Scientific, Waltham, MA, USA) equipped with heated electrospray ionization source (capillary temperature 360°C, sheath gas 60 nominal units, aux gas 20 nominal units, sweep gas 5 nominal units, aux gas temperature 400°C, spray voltage 3.3 kV, S-lens RF level 50) operated in positive or negative ion mode. Full scan measurements (*m*/*z* 100-1200, resolution 280k, AGC target 3 . 10^6^, maxIT 900 ms) were performed separately for positive and negative ion mode. MS^2^ scans with metabolite-specific fragmentation energies were used for metabolite identification.

### Isotopologue detection

Full scan RAW files were converted to mzXML using ProteoWizard msConvert (53) with the vendor’s algorithm for peak picking. Isotopologue detection was achieved with R-based X^13^CMS (27). The R script can be found in the Supplementary Information. In brief, after peak-picking with centwave (3 ppp, peakwidth 5-20 s) and retention time alignment with orbiwarp, isotopologues with a mass difference of 1.00335 Da were assigned (RTwindow 10 s, 3 ppm). Either all isotopologue groups (α = 1) or only isotopologue groups significantly different from ^12^C (α = 0.05) were reported. Afterwards, significantly increased isotopologue groups were manually curated with reference to the original spectrum in order to exclude groups that did not contain at least 3 consecutive isotopologues.

### Compound identification

Compounds were identified based on their retention time, high resolution mass to charge ratio and fragmentation pattern. Compound Discoverer (Vers. 2.1, Thermo Fisher Scientific, Waltham, MA, USA) was used to predict sum formula, search an in-house and public databases (ChemSpider and mzCloud) as well as calculate FISh scores of candidates. SIRIUS and CSI:FingerID were used to compute fragmentation trees and search molecular structure databases (54). Putatively identified compounds were compared to authentic standards: Arachidonic acid, betaine, carnitine hydrochloride, choline chloride, 11*Z*-eicosenoic acid, 2-hydroxytetradecanoic acid, isoleucine, leucine and 1-oleoyl-*sn*-glycero-3-phosphocholine were obtained from Sigma Aldrich (Munich, Germany). Oleic acid was purchased from AppliChem (Darmstadt, Germany). Linoleic acid was from Alfa Aesar (Haverhill, MA, USA). 5*Z*,8*Z*,11*Z*,14*Z*,17*Z*-Eicosapentaenoic acid was supplied by Cayman Chemicals (Ann Arbor, MI, USA) and 6*Z*,9*Z*,12*Z*-Hexadecatrienoic acid from Larodan (Solna, Sweden). 6*Z*,9*Z*,12*Z*,15-Hexadecatetraenoic acid has been synthesized according to Pohnert, Adolph and Wichard (45). ß-Alanine betaine has been synthesized according to Chary, Kumar, Vairamani and Prabhakar (55) Leucine and isoleucine were not baseline separated and are consequently grouped as (iso-)leucine.

### Calculation of the degree of labeling

To calculate the degree of labeling isotopologues are assumed to have a Bernoulli distribution (see Supplementary Information for formula).(56) Measured isotopologue intensities are compared to computed distributions and the squared coefficient of variation between both is minimized in an iterative process.

## Acknowledgments

We thank W. Kooistra for providing diatom cultures. David Russo is acknowledged for helpful comments on an initial draft of this manuscript.

## Author contributions

N.M. developed the pipeline and analyzed the data. A.R. acquired and analyzed data. N.M. and G.P. conceived the study, directed all experiments, and wrote the manuscript with contributions from the co-author. All authors approved the manuscript.

## Data and materials availability

All mass spectra are deposited in the EMBL metabolomics data repository Metabolights. All computer code (in R) developed for this study is available in the supplementary information.

## Supporting Information

This article contains supporting information online at…

## Supplementary Materials

**Fig. S1.**
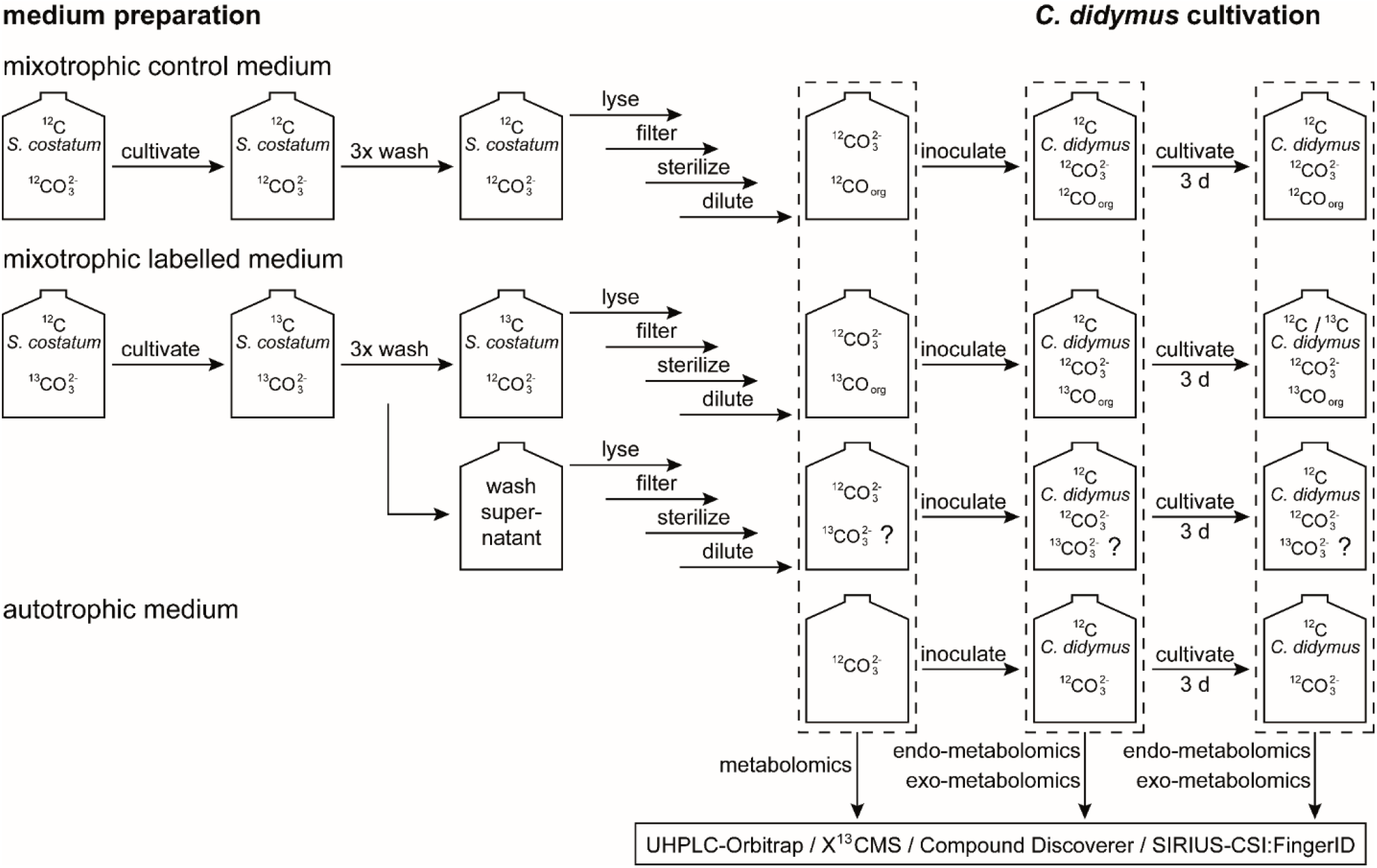
Experimental setup for mixotrophy experiment. Last two lines: To demonstrate the effective removal of inorganic ^13^C, *C. didymus* was grown on the wash supernatant and did not contain labelled metabolites

**Fig. S2.**
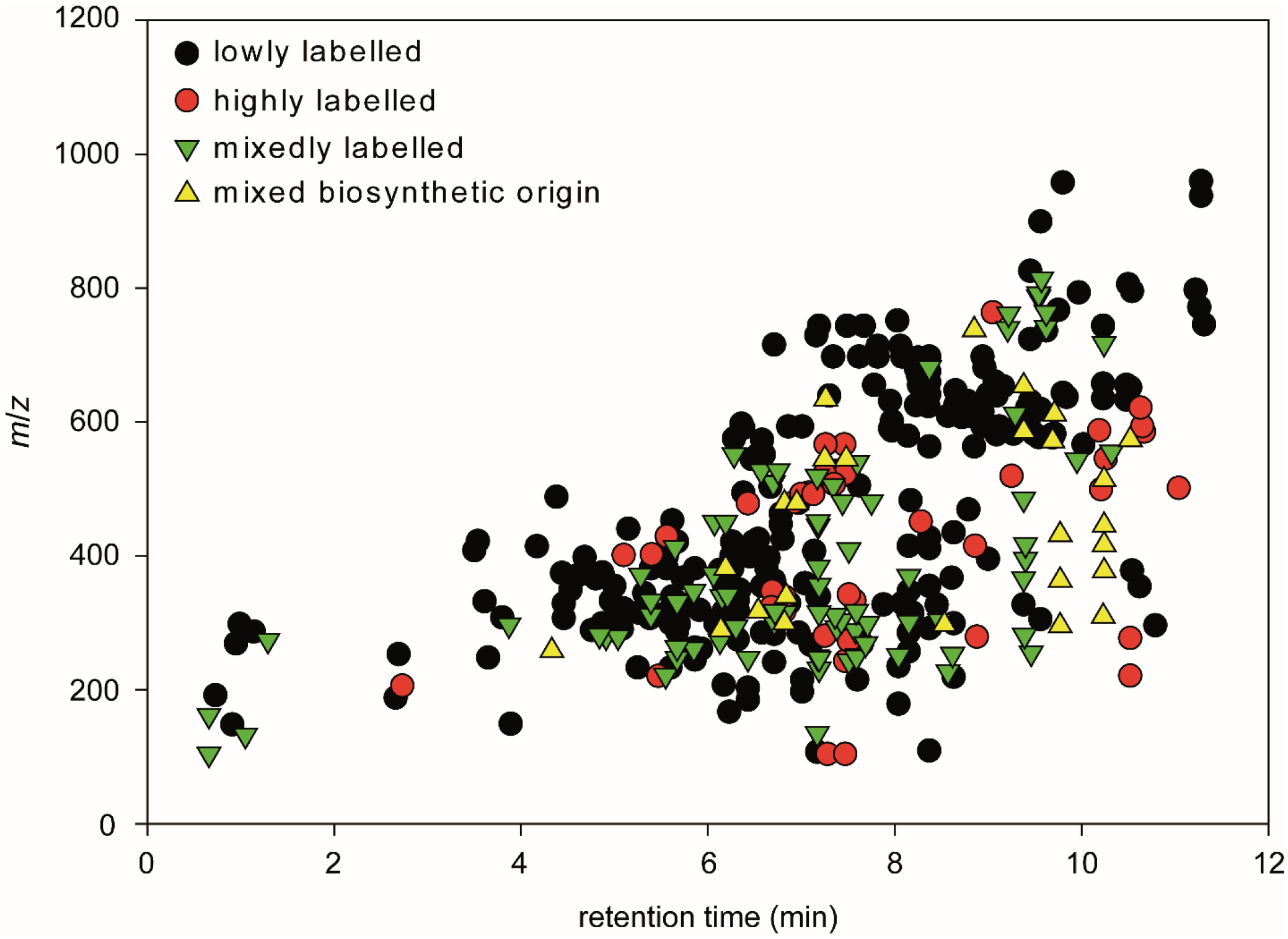
Labelling of metabolites in *Chaetoceros didymus* endometabolome. Correlation between retention time (gradient as described in materials and methods) and *m*/*z* of manually curated isotopologue groups sorted by labelling pattern as described in Fig. 1.

**Fig. S3.**
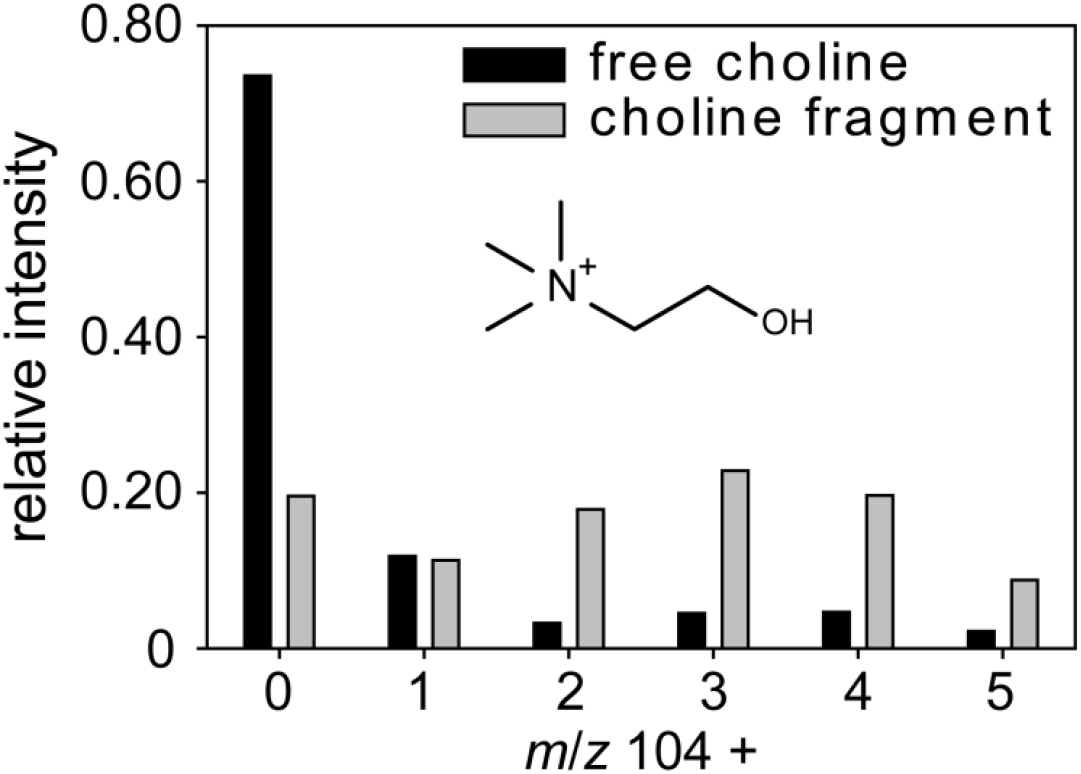
Labelling pattern of carnitine. Depicted are the isotopologues of carnitine. The presence of M+5 to M+7 proves labelled *N*-methyl groups.

## R-script for X^13^CMS analysis

~~~
require(xcms)
require(X13CMS)
# set working directory to one where the “C12” and “C13” folders reside
setwd(“E:/X13CMS”)
# Peak-picking and retention-time alignment with XCMS
xs= xcmsSet(c(‘./C12′, ‘./C13′), method= ‘centWave’, ppm= 3, peakwidth= c(5, 20))
xs= group(xs, bw=5, mzwid=0.015)
xs2= retcor(xs, method= ‘obiwarp’)
xs2= group(xs2, bw=5, mzwid=0.025)
xs3= fillPeaks(xs2)
# Setting variables for X13CMS
sN = rownames(xs3@phenoData) # sample names
sN = sN[c(1:3, 4:6)] # samples (3 unlabeled, 3 labeled)
# -----only significantly different isotopologues ------
# labeling report for samples:
labelsSign = getIsoLabelReport(xcmsSet = xs3, sampleNames = sN, unlabeledSamples = “C12”, labeledSamples = “C13”,
isotopeMassDiff = 1.00335, RTwindow = 10, ppm = 3, massOfLabeledAtom = 12, noiseCutoff = 10000, intChoice =
“intb”, varEq = FALSE, alpha = 0.05, singleSample = FALSE, compareOnlyDistros = FALSE, monotonicityTol = FALSE,
enrichTol = 0.1)
# in each of the sN variables, the first 3 samples listed are of the “C12” or unlabeled type while the next 3 are of the “C13”
type
classes = c(rep(“C12”,3), rep(“C13”,3))
# print labeling report to a text file (recommended to open in Excel)
printIsoListOutputs(listReport = labelsAll, outputfile = “all/labels_all.txt”)
# print pdf of isotopologue groups in a single labeling report plotted as relative intensity distributions
plotLabelReport(isoLabelReport = labelsAll, intOption = “rel”, classes, labeledSamples = “C13”, outputfile =
“all/labelsrel_all.pdf”)
# print pdf of isotopologue groups in a single labeling report plotted as absolute intensity distributions
plotLabelReport(isoLabelReport = labelsAll, intOption = “abs”, classes, labeledSamples = “C13”, outputfile =
“all/labelsabs_all.pdf”)
~~~

## Bernoulli statistics to calculate the degree of labelling

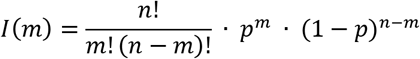

For a metabolite with *n* carbon atoms the intensity of an isotopologue *I(m)* with *m* 13C atoms is calculated using the degree of labelling *p*.

